# Genomic epidemiology and strain taxonomy of *Corynebacterium diphtheriae*

**DOI:** 10.1101/2021.07.18.452871

**Authors:** Julien Guglielmini, Melanie Hennart, Edgar Badell, Julie Toubiana, Alexis Criscuolo, Sylvain Brisse

## Abstract

**Background:** *Corynebacterium diphtheriae* is highly transmissible and can cause large diphtheria outbreaks where vaccination coverage is insufficient. Sporadic cases or small clusters are observed in high-vaccination settings. The phylogeography and short timescale evolution of *C. diphtheriae* are not well understood, in part due to a lack of harmonized analytical approaches of genomic surveillance and strain tracking.

**Methods:** We combined 1,305 genes with highly reproducible allele calls into a core genome multilocus sequence typing (cgMLST) scheme. We analyzed cgMLST genes diversity among 602 isolates from sporadic clinical cases, small clusters or large outbreaks. We defined sublineages based on the phylogenetic structure within *C. diphtheriae* and strains based on the highest number of cgMLST mismatches within documented outbreaks. We performed time-scaled phylogenetic analyses of major sublineages.

**Results:** The cgMLST scheme showed high allele call rate in *C. diphtheriae* and the closely related species *C. belfantii* and *C. rouxii*. We demonstrate its utility to delineate epidemiological case clusters and outbreaks using a 25 mismatches threshold, and reveal a number of cryptic transmission chains, most of which are geographically restricted to one or a few adjacent countries. Subcultures of the vaccine strain PW8 differed by up to 20 cgMLST mismatches. Phylogenetic analyses revealed short timescale evolutionary gain or loss of the diphtheria toxin and biovar-associated genes. We devised a genomic taxonomy of strains and deeper sublineages (defined using a 500 cgMLST mismatches threshold), currently comprising 151 sublineages, only a few of which are geographically widespread based on current sampling. The cgMLST genotyping tool and nomenclature was made publicly accessible at https://bigsdb.pasteur.fr/diphtheria.

**Conclusions:** Standardized genome-scale strain genotyping will help tracing transmission and geographic spread of *C. diphtheriae*. The unified genomic taxonomy of *C. diphtheriae* strains provides a common language for studies into the ecology, evolution and virulence heterogeneity among *C. diphtheriae* sublineages.

## Introduction

*Corynebacterium diphtheriae* causes diphtheria, a severe infection of the upper respiratory tract. The most typical symptoms include the formation of pseudomembranes in the throat, and toxin-induced complications such as myocarditis and neuropathy. The diphtheria toxin is produced by some strains that carry the *tox* gene, harbored by a lysogenized corynephage. Cutaneous infections by *C. diphtheriae* are common and skin carriage or infection may act as source of respiratory infections [1]. Isolates without *tox* gene can cause invasive infections such as endocarditis and sepsis [2]. *C. diphtheriae* is considered to spread predominantly through droplets among humans, and is only rarely isolated from animals. Strains of the phylogenetically related species *C. ulcerans* and *C. pseudotuberculosis* can also produce the diphtheria toxin, but in these cases human infections always result from transmission from animals.

Diphtheria is largely eliminated in countries with high coverage rates of vaccination with the diphtheria toxoid [3]. Nevertheless, the pathogen *C. diphtheriae* continues to circulate among humans and even in high-vaccination coverage countries, sporadic cases of toxigenic *C. diphtheriae* infections are reported regularly. In Europe, 139 cases were reported between 2014 and 2018, a majority of which followed introduction of *C. diphtheriae* from endemic regions [4]. In these settings, transmission is rare and only small clusters of patients or carriers are observed [5, 6]. Differently, in settings where public health action is disrupted and vaccination coverage has decreased, several large outbreaks of diphtheria in its typical clinical form have occurred in recent years [7–9].

Strain subtyping is used to study genotype-phenotype relationships and to define chains of transmission and the origin of outbreak strains. The subdivision of *C. diphtheriae* into four biovars is of poor resolution and has no epidemiological usefulness, and even its relevance for strain classification is questionable [10, 11]. The recently described species *C. belfantii* [12] and *C. rouxii* [13] include most strains that were previously assigned to *C. diphtheriae* biovar Belfanti. Although no toxigenic strains of these two novel species were described so far, they are typically misidentified as *C. diphtheriae* when using reference molecular tests, biochemical diagnostic methods and even MALDI-TOF mass spectrometry [12, 14][13].

The highly standardized molecular methods ribotyping [15] and 7-gene MLST [14] have been developed to subtype *C. diphtheriae*, but in the genomic era their resolution power appears to be highly limited [11]. Whole genome sequencing (WGS) provides nearly ultimate resolutive power, and several recent studies have used this approach for *C. diphtheriae* population biology [10, 16–18] or molecular epidemiology and outbreak investigation [5, 19–22][23]. However, currently a shared genotyping system and strain definition standard are lacking, which limits our ability to compare strains across studies.

Among WGS analytical approaches, cgMLST [24] has several attractive characteristics, including high reproducibility and portability that allow strain matching across studies, geography and time. cgMLST has been used successfully to define transmission of *C. diphtheriae* among patients [5, 20, 21]. However so far, no cgMLST scheme is provided through a publicly available web application.

The aim of this work was to address this gap by providing an accessible genotyping standard and harmonized definitions of strains and sublineages. We calibrated the discriminatory power of cgMLST by comparing this proposed standard to whole genome single nucleotide polymorphisms (SNPs), and illustrate how cgMLST can be used to identify strains from recent *C. diphtheriae* clusters and outbreaks. Last, we mapped 7-gene MLST sequence types (ST) identifiers to the novel cgMLST-based sublineage nomenclature, which provides backwards compatibility with this previous classification framework.

## Material and Methods

### Isolates and outbreak sets included in the study

A total of 440 isolates (**Table S1**) were included into the study dataset. First, we included and sequenced 8 isolates of the French National Reference Center (NRC) collection corresponding to four available pairs of *C. diphtheriae* infection cases defined as epidemiologically linked (**Table 1**). The first pair (Mayotte set 1) included two isolates from a single sample, isolated at one-month distance (FRC025 and FRC030). The second pair (Mayotte set 2) included isolates from one patient and one of his contacts; the two isolates were isolated 15 days apart. The third pair (Carcassonne set 1) included isolates from two members of the same household, sampled 14 days apart. The fourth pair (Bron set 1) included two isolates from a pair of siblings, sampled on the same day. Second, we included and sequenced a random selection of 21 isolates from the French NRC collection, selected among isolates that were co-circulating temporally or geographically with each outbreak (**Table S1,** ‘sporadic’ with FRCxxxx isolate names). Third, 162 isolates from previously published genomic epidemiology studies in South Africa [25], Germany-Switzerland [5], Germany [20], France [26] and Canada [21] were included. These genomic investigations of outbreaks (or clusters of cases) together comprised 11 sets of isolates defined as related based on epidemiologically and genomic evidence (**Table 1**), and also included isolates considered as non-related to the outbreak sets. Finally, all other publicly available genomic sequences from GenBank available on May 30^th^ 2019 were included (**Table S1**). Of these, we removed isolates from the Canadian study (isolates CD46 to CD49 and CD51 to CD56) that were of low quality. We also removed 3 isolates with atypical assemblies: 2 from Germany (KL0932 and KL0954) because they were highly fragmented (446 and 496 contigs, respectively) and one from a Germany-Switzerland study (Cd_14) because of a low G+C content (42.9%).

**Table 1.** Outbreak datasets.

| Dataset | No. of strains | 7-gene MLST | Sublineage | Genetic cluster | Country | Year | Outbreak information | Reference |
| --- | --- | --- | --- | --- | --- | --- | --- | --- |
| South Africa set 1 | 19 | 378 | 378 | 472 | South Africa | 2015-2016 | tox-positive | du Plessis *et al.*, 2017 |
| South Africa ST395 | 5 | 395 | 395 | 68;491;492;542 | South Africa | 1980;2015 | tox-negative | du Plessis *et al.*, 2017 |
| Germany-Switzerland set 1 | 3 | 473 | 473 | 13 | Germany | 2015 | Refugees from Eritrea, Ethiopia, Somalia and Syria; tox-positive | Meinel *et al.*, 2016 |
| Germany-Switzerland set 2 | 3 | 183;486 | 486 | 204 | Germany | 2015 | Refugees from Eritrea, Ethiopia, Somalia and Syria; tox-positive | Meinel *et al.*, 2016 |
| Germany-Switzerland set 3 | 4 | 398 | 398 | 207 | Germany | 2015 | Refugees from Eritrea, Ethiopia, Somalia and Syria | Meinel *et al.*, 2016 |
| Germany cluster 1 | 14 | 8 | 8 | 414 | Germany | 2016-2017 | Hamburg, Essen, Kiel | Dangel *et al.*, 2018 |
| Germany cluster 2 | 19 | 8 | 8 | 292 | Germany | 2015-2017 | Berlin mostly | Dangel *et al.*, 2018 |
| Germany cluster 3 | 6 | 8 | 8 | 411 | Germany | 2015-2016 | Berlin, Essen, Trier | Dangel *et al.*, 2018 |
| Germany cluster 4 | 2 | 8 | 8 | 415 | Germany | 2016-2017 | Hannover | Dangel *et al.*, 2018 |
| Germany ST130 | 10 | 130 | 130 | 402 | Germany | 2012-2017 | None | Dangel *et al.*, 2018 |
| Vancouver cluster 1 | 43 | 76;746 | 76 | 456 | Canada | 2017-2018 | Vancouver inner city, cutaneous infection cluster | Chorlton et al., 2019 |
| Mayotte set 1 | 3 | 97 | 97 | 78 | Mayotte | 2008-2009 | Single sample, 1 month apart | This study |
| Mayotte set 2 | 2 | 184 | 184 | 86 | Mayotte | 2009 | Patient + contact, 15 days apart | This study |
| Carcassonne set 1 | 2 | 212 | 212 | 222 | France | 2011 | Members of household, 14 days apart | This study |
| Bron set 1 | 2 | 411 | 411 | 116 | France | 2016 | Siblings, sampled on same day | This study |

While this work was in progress, additional *C. diphtheriae* genomes were published from Austria [27], the USA [23], Yemen [9], France [11], Spain [28] and India [29]. We included these 162 genomes in a joint phylogenetic analysis with the 440 above genomes (**Figure Suppl. 1**).

### DNA preparation and sequencing

Isolates were retrieved from −80°C storage and plated on tryptose-casein soy agar for 24-48 hours. A small amount of bacterial colony biomass was re-suspended in a lysis solution (Tris-HCl pH8, 20mM, EDTA mM, Triton X-100 1.2% and lysozyme 20mg/ml) and incubated at 37°C for 1 h and DNA was extracted with the DNeasy blood&Tissue kit (Qiagen, Courtaboeuf, France) following the manufacturer’s instructions. Genomic sequencing was performed using a NextSeq500 instrument (Illumina, San Diego, USA) with a 2 x 150 nt paired-end protocol following Nextera XT library preparation [11].

### Genomic sequence assembly

For *de novo* assembly, paired-end reads were clipped and trimmed using AlienTrimmer v0.4.0 [30], corrected using Musket v1.1 [31] and merged (if needed) using FLASH v1.2.11 [32]. For each sample, remaining processed reads were assembled and scaffolded using SPAdes v3.12.0 [33].

### Definition of the cgMLST scheme

We first selected 171 genomes representing a diversity of *C. diphtheriae* sublineages, including 104 genomes present in **Table S1** and 67 genomes from the French NRC [11]. These included 14 *C. belfantii* isolates, but no *C. rouxii* isolates were included at this step, as this species was not identified at that time. From this set of genomes, we inferred the species core genome using the CoreGeneBuilder pipeline (https://zenodo.org/record/165206#.WVpT7I55EQU) as in [34], using the NCTC13129 strain genome (GenBank accession no. NC_002935) as reference [35]. CoreGeneBuilder automatically removed genomes for which the size was too divergent compared with the whole set. We applied the following quality filters: a maximum of 500 contigs and a minimum N50 of 10,000 bp. With those criteria, no genome was filtered out. The pipeline’s next step relies on eCAMBer [36], which consists of a *de novo* annotation of the genomes (except the reference) using Prodigal [37] and in the harmonization of the positions of the stop and start codons. In the last step, the core genome is inferred using a bidirectional best hits approach, following Touchon et al. [38]. We used CoreGeneBuilder default settings and considered a gene as part of the core genome if it was found in at least 95% of the 171 selected genomes. This resulted in an initial core genome containing 1,555 loci.

To obtain a set of loci that would be highly robust to genotyping artifacts, we filtered out some genes based on several criteria. First, we removed potential paralogs, as the presence of paralogs inside a typing scheme can lead to genotyping errors when a gene sequence is attributed to the wrong locus. To detect those potential paralogs, we compared each allele of each locus against all the alleles of all the other loci using the software BLAT [39]. If a single hit was found between 2 different loci (more than 70% amino acid sequence identity between two alleles), we removed both. A total of 8 loci were discarded by this process. We also removed genes that belong to the classical 7-gene MLST scheme [14] and the ribosomal protein genes that are used in the ribosomal MLST approach [40], so that they could be analyzed independently. Next, we removed loci whose length varies too much among alleles, which is useful to reduce ambiguities during the genotyping process. We aligned the amino acid sequences and removed loci for which the alignment contained more than 5% of gaps (total number of gaps compared with the total number of character states). This filter removed 107 loci. In addition, we removed 3 loci because at least 1 allele showed 1 or more ambiguous character state(s). Altogether, these filters resulted in a list of 1388 remaining loci.

To filter out loci with non-reproducible allele calls at low assembly coverage depth, randomly selected read pairs representing coverage depths of 10X to 50X were drawn (five times for each isolate/coverage combination) from the raw sequencing data of 4 isolates (*C. diphtheriae* 18-093a, FRC0024; and *C. belfantii* FRC0019 and FRC0318); we included two *C. belfantii* isolates because their genome is more fragmented due to a high copy number of insertion sequences [41]. After assembly and allele calling using BIGSdb, we eliminated 33 loci that showed non-reproducible allele calls from all assemblies obtained with coverage depths equal or above 20X in any of the four isolates.

Finally, we also removed 52 loci that were uncalled in >10% of scanned genomes. These filtering steps led to a final set of 1,305 gene loci, which together constitute the ‘Pasteur *C. diphtheriae* cgMLST scheme’, or hereafter ‘cgMLST scheme’ in short. Note that a given gene locus could have been filtered out by several filters applied independently, so that the sum of numbers of loci removed by individual filters is smaller than the final number of loci that were filtered out.

### Evaluation of cgMLST allele calling rate

Because loci with high numbers of missing alleles were filtered out during cgMLST scheme construction, the allele call rate of the resulting scheme was assessed using an independent set of genomes, which had not been used for the above-described definition of the scheme. This entirely distinct dataset of *C. diphtheriae* isolates was made of 162 genomes published after cgMLST scheme definition and described in the first Methods paragraph. Missing alleles were defined as being not called using 90% identity and 90% of length coverage.

### Constitution of a BIGSdb database for *C. diphtheriae*

The cgMLST scheme was incorporated into a BIGSdb seqdef database [42]. Classification schemes were created at the level of 25 and 500 mismatches. The isolates provenance data and genome sequences were imported into a linked isolates database (https://bigsdb.pasteur.fr/diphtheria). A publicly available BIGSdb project (*i.e*., a browsable list of isolates) entitled ‘cgMLST set-up study’ corresponds to the dataset of 440 genomes and was created to facilitate retrieval of the data used in this work.

### Inheritance of classification identifiers from 7-gene MLST

MLST alleles and profiles of the 440 genomes were defined in the PubMLST *C. diphtheriae* database. We then mapped the sequence type (ST) identifiers onto the obtained 500-mismatch cgMLST single linkage classification groups, and attributed the dominant ST identifier to the group, using an inheritance algorithm (https://gitlab.pasteur.fr/BEBP/inheritance-algorithm).

### Comparison with other schemes

An ad-hoc cgMLST scheme for *C. diphtheriae* was defined and used previously [19, 20, 43]. In order to compare our scheme with this previous one, we took the first allele of all loci of both schemes and performed an all-against-all BLASTN analysis. Hits were defined as sequence pairs with a proportion of identical nucleotides greater than 90% and with an alignment length longer than half of both the query and the subject sequences.

### Cryptic cluster detection

Cryptic clusters were defined as single-linkage groups of genomes (i) defined using a threshold of 25 cgMLST allelic mismatches, but (ii) which were not initially defined as clusters based on available epidemiological data.

### Phylogenetic and population structure analyses

Phylogenetic analyses were performed by first inferring a tree using JolyTree, an alignment-free distance-based procedure [44]. The resulting tree was used as input for ClonalFrameML [45], which also requires an alignment. The alignment was obtained by translating allelic sequences of cgMLST loci of each genome into proteins. We aligned the proteins of each loci using MAFFT v7.467 [46] with default parameters, and then back-translated the protein alignments to codon alignments using Goalign (https://github.com/evolbioinfo/goalign); the individual gene alignments were then concatenated into a supermatrix, also using Goalign (the nucleotide positions of missing alleles were replaced by gaps in the concatenate of alignments of individual loci). We used this alignment and above defined input tree to run ClonalFrameML with default parameters. We used the software PHIPack v1.0 [47] to calculate the PHI (Pairwise Homoplasy Index) statistic and perform a locus-by-locus test of recombination with a *p*-value threshold of 0.05. Nucleotide diversity was calculated both for the entire dataset of each species, and among sublineages, using an in-house script relying on the *p*-distance calculation provided by Goalign. The overall average Silhouette width, which estimates the overall clustering quality based on maximizing inter-group distances and minimizing intra-group distances, was used to define the clustering consistency [48].

Dating ST5 and ST8 phylogenetic tree nodes was performed using the LSD (Least-Squares Dating) algorithm [49] as implemented in IQ-TREE v2.0.6 [50]. The sampling dates were used to calibrate the trees. The subtrees were extracted from the entire tree (**Figure 1**). For both analyses, the closest outgroup was also extracted and included in the analysis. This allowed for the dating of the root of the ST8 phylogeny, whereas for ST5 this analysis did not allow dating the root.

**Figure 1.**
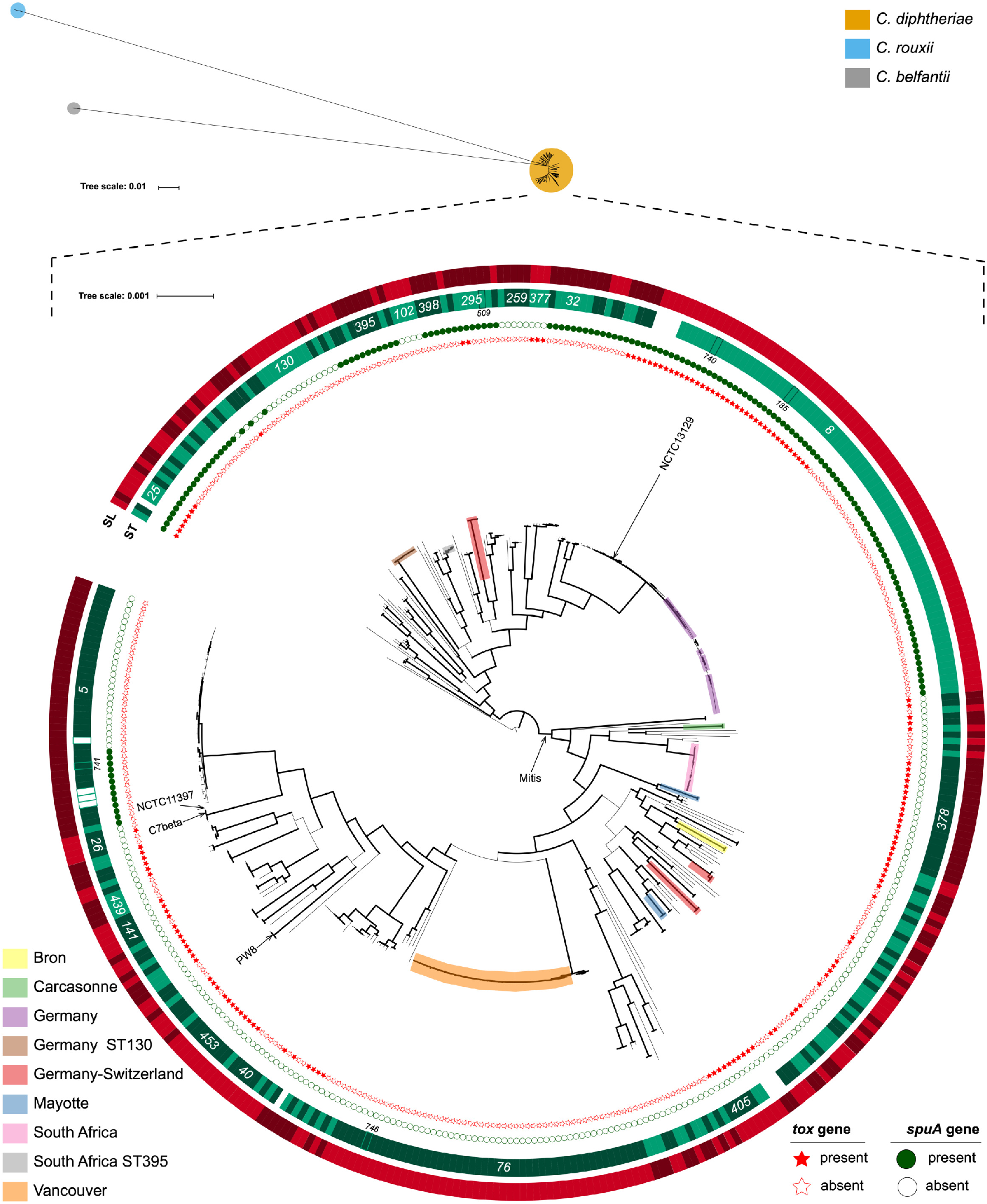
Phylogenetic tree of *C. diphtheriae* sublineages. Maximum-likelihood phylogenetic tree for 440 *C. diphtheriae, C. belfantii and C. rouxii* isolates based on the concatenated multiple sequence alignments of 1,305 core genome multilocus sequence typing loci. The top panel represents the unrooted topology of the three species. The bottom panel is the *C. diphtheriae* subtree. Branches with support values greater than 70% are thicker. The root of this subtree corresponds to the placement of the branch leading to *C. rouxii* and *C. belfantii* in the top panel tree. The *C. diphtheriae* subtree contains the following annotations: The stars represent the presence (red star) or absence (white star) of the diphtheria toxin *tox* gene. The small circles represent the presence (green circle) / absence (white circle) of the *spuA* gene. The green strips represent the alternation of 7-gene MLST STs of the isolates: when the ST changes while walking along the circle, the shade of green also changes. Identifiers of the main STs are indicated. When an ST could not be attributed (because at least one of the 7 genes was missing), the color is white. A strip with a visible border represents isolates with an ST that is different from the rest of the clade, and the ST number is provided below. For example, ST185 is found within the ST8 clade. The red strips follow the same alternation logic for the cgMLST sublineages identifiers. Note the strong concordance between ST and cgMLST sublineages. The position of noteworthy reference isolates is given by black arrows. Finally, the epidemic clusters’ positions are indicated by colored boxes on the corresponding branches (see bottom left key).

The clade compatibility index of STs or other groups was calculated using the ETE Python library (http://etetoolkit.org/docs/latest/tutorial/tutorial_trees.html#checking-the-monophyly-of-attributes-within-a-tree), in order to define whether their constitutive genomes formed a monophyletic, paraphyletic or polyphyletic group within the recombination-purged sequence-based phylogeny of the core genome. We estimated clade compatibility as the proportion of non-singleton STs, sublineages or clonal groups that were monophyletic.

### Whole-genome SNP calling based on a read mapping approach

We used Snippy v4.4.0 [51] with default parameters to perform SNP calling within the German and Canadian outbreak datasets, using as references, strains NCTC13129 and CD50 (from the BioProject PRJNA563223), respectively.

### Saturation analysis of the number of sublineages and genetic clusters

Following the definitions of genetic clusters and sublineages (25 allelic mismatches for the genetic clusters, 500 for the sub-lineages), we simulated 1,000 samples of 1 to 440 profiles (the 440 profiles of our initial dataset), with a step of 1. For each threshold and each number of profiles, we then calculated the mean number of clusters, and used the minimum and maximum values to define the range. To extrapolate the previously-obtained curves, we defined the species accumulation curves using the R package *vegan*. We chose the Lomolino function [52] for its excellent fit with our data.

## Results

### Definition and evaluation of a cgMLST scheme for genotyping isolates of *C. diphtheriae, C. belfantii* and *C. rouxii*

We defined a set of 1,305 protein-coding gene loci deemed appropriate for *C. diphtheriae, C. belfantii* and *C. rouxii* genotyping (see Methods). These loci were combined into a cgMLST scheme and altogether have a total length of 1,265,478 nucleotides (calculated for allele 1 of each gene), representing 51% of the genome size of the ST8 reference strain NCTC13129. Individual locus lengths varied from 93 to 4008 nt. The list of loci and their characteristics are presented in **Table S2.**

The allele call rate of the cgMLST scheme loci was determined using posteriorly published genomes from Austria [27], the USA [23], Yemen [9], France [11], Spain [28] and India [29] and was found to be 98.6% ± 1.5%. We also calculated the call rate using 337 additional, prospectively collected genome sequences from the French NRC for diphtheria (unpublished). The mean number of called alleles was 1,288 (98.7%) and 1,287 (98.6%) for *C. diphtheriae* and *C. belfantii*, respectively. We noted that a few *C. belfantii* genomes had a lower call rate (**Figure S2**). For *C. rouxii*, allele call rate was lower (1,190; 91.2%), as expected since this novel species was not included during scheme development and is genetically more distant [53]. The allele call rate was not significantly different between *tox*-positive and *tox*-negative *C. diphtheriae* (98.8 versus 98.6%, respectively; **Figure S3**). The relationships between allele call rate and genome assembly fragmentation showed that high call rates (>95%) were generally obtained, even for *C. diphtheriae* genome assemblies that were fragmented into >400 contigs (**Figure S2**). Overall, these results demonstrate high typeability across the phylogenetic breadth of *C. diphtheriae* and its two closely related species *C. belfantii* and *C. rouxii*.

Based on the study dataset of 440 genomes, allele numbers ranged from 7 to 285 per locus, illustrating the high amounts of genetic diversity of the core genes within *C. diphtheriae*. As expected, the number of alleles per locus increased with locus size (**Figure S4**). Locus-by-locus recombination analyses detected evidence of intra-gene recombination within a majority of loci (843; 64.6%), whereas only 431 (33%) loci were deemed non recombinogenic. There were 31 (2.4%) loci with too few polymorphic sites to be tested (**Table S2**). The number of alleles per locus was higher for recombinogenic loci (**Figure S5**), which reflects the contribution of homologous recombination to the generation of allelic diversity.

We compared the cgMLST scheme defined herein (named the Pasteur scheme) to a previously published *ad-hoc* (not publicly available for comparison) cgMLST scheme [20], and found 1132 loci in common, representing 87% of the Pasteur cgMLST loci (**Figure S6**). Of these, 115 varied slightly in length due to alternative start codon definitions. Whereas 173 loci were unique to the Pasteur scheme, 421 were unique to the previous scheme. Three loci were not counted as common to the two schemes even though their sequences matched, because they are encoded on opposite strands.

### Population structure of *C. diphtheriae*

A phylogenetic analysis was conducted based on a recombination-purged concatenate of the individual alignments of the 1,305 loci (**Figure 1**). The three deepest branches corresponded to the three species *C. rouxii*, *C. belfantii* and *C. diphtheriae*. Whereas *C. diphtheriae* included most isolates (86.5%, 148/171), *C. rouxii* and *C. belfantii* comprised 6 and 17 isolates, respectively. MLST data (**Figure S7**) revealed 134, 10 and 3 STs within *C. diphtheriae, C. belfantii* and *C. rouxii*, respectively. The nucleotide diversity, estimated as the average SNP distance among isolates, was 0.012 for *C. diphtheriae*, 0.0025 for *C. belfantii* and 0.0039 for *C. rouxii*. The average genetic distance between sublineages (*i.e*., considering only one representative genome of each sublineage, as defined below) was 0.024 polymorphism per nucleotide position.

The phylogenetic structure within *C. diphtheriae* revealed a star-like phylogeny with multiple deeply-branching sublineages as previously reported [11, 18]. The phylogenetic distribution of 7-gene MLST data showed that sublineages typically corresponded to a single ST (**Figure 1**). Some sublineages comprised closely-related isolates, likely to be the result of sampling from case clusters, outbreaks or recent clonal expansions (**Table S1**).

Diphtheria toxin gene *tox*-positive isolates were observed in 19 sublineages distributed on the Southern hemisphere of the tree presented in **Figure 1**, including the sublineage that includes the vaccine strain PW8, and in 5 sublineages in the Northern hemisphere of the tree, including the one that comprised strain NCTC13129 [11]. These two hemispheres of the tree are identifiable by the distribution of the *spuA* gene, involved in starch utilization [54]: the upper lineage was *spuA*-positive with the exception of six *spuA*-negative sublineages, whereas in sharp contrast, the lower lineage was almost entirely *spuA* negative, with ST5 being the single exceptional *spuA*-positive sublineage. The two major lineages of *C. diphtheriae* were therefore labeled as lineage Gravis and Mitis, respectively [11]. The addition of 162 recently published genomes (**Figure S1**) fully confirmed the above observations. In addition to ST5, sublineage ST466 belonged to the lineage Mitis and included *spuA*-positive isolates. These *spuA*-positive isolates were closely related to two *spuA* negative isolates, and all ST466 genomes were from India, suggesting a recent acquisition in this country of the *spuA* gene, along with probable conversion of the biovar from Mitis to Gravis.

### Definition of *C. diphtheriae* sublineages and their geographic distribution

To guide the definition of *C. diphtheriae* sublineages, we performed an analysis of the mathematical properties of the clustering of cgMLST profiles at all possible thresholds (Silhouette analysis; **Figure S8**). We observed optimal clustering indices on a large interval (~between 300 and 600 mismatches) with a maximum value at 554. As the Silhouette index was almost maximum at 500 cgMLST mismatches, for simplicity we chose this value to define *C. diphtheriae* sublineages (SL). Our dataset of 440 genomes comprised 114 sublineages, and a saturation curve analysis (**Figure 2**) indicates that future addition of *C. diphtheriae* genomes is expected to reveal the existence of a multitude of additional sublineages that remain to be sampled. The additional dataset of 162 genomes uncovered 22 additional sublineages (**Figure S1**). The average core genome nucleotide divergence among sublineages was 2.4%.

**Figure 2.**
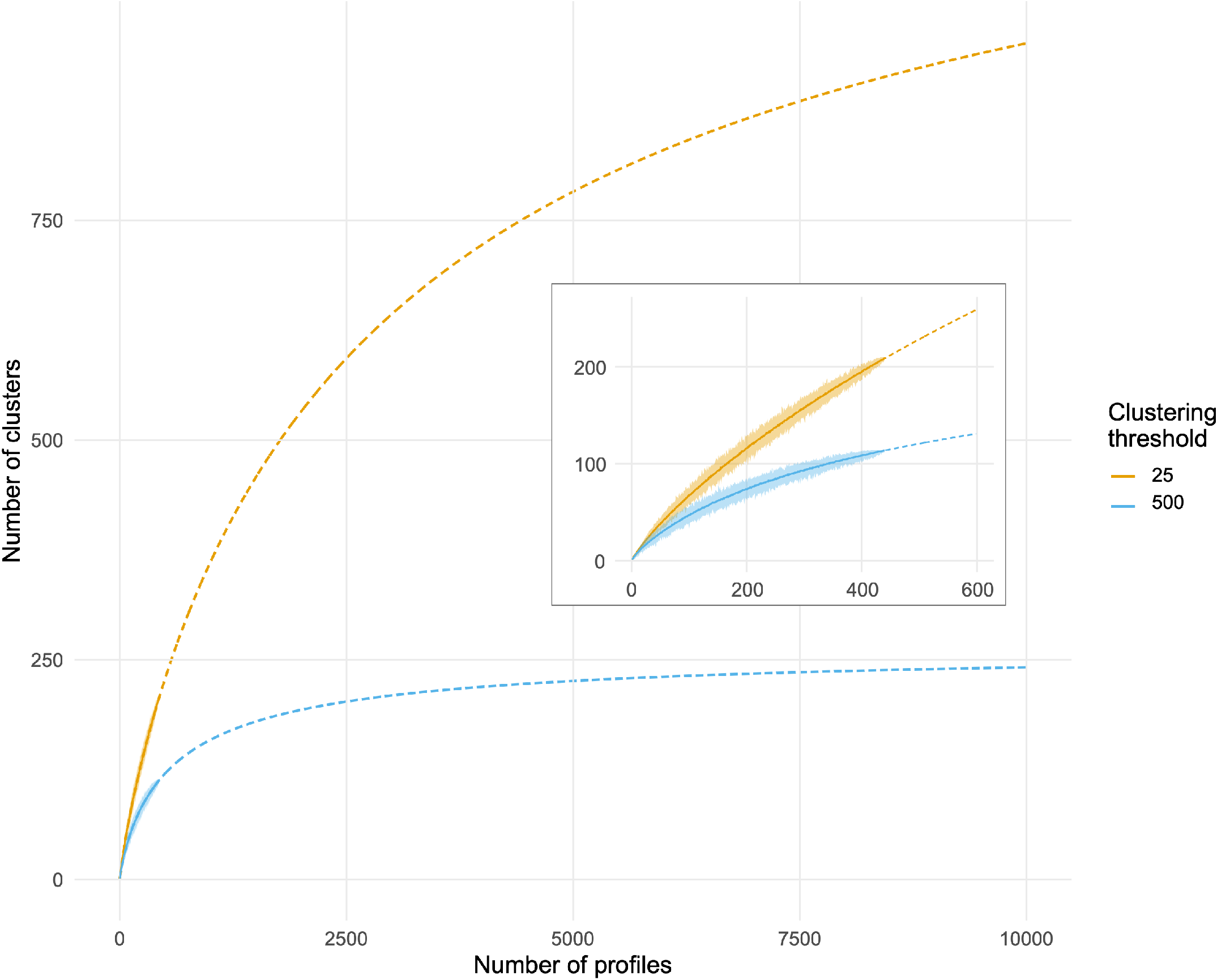
Saturation analysis of the number of sublineages and genetic clusters. The curves in the inset correspond to the mean number of clusters found of each clustering threshold, and the area around the curves represents the range (min to max). The main panel represents the extrapolation (see Methods).

To create continuity in the numerical identifiers of sublineages with respect to MLST, we mapped the ST nomenclature on sublineages and incorporated the resulting sublineage tags into the BIGSdb *C. diphtheriae* genotype nomenclature database (**Table S3**). Note that there was a strong concordance between MLST and cgMLST sublineages (**Figure S9**) and that this correspondence was established up to ST750 (November 17^th^ 2020).

All non-singleton cgMLST sublineages (66/66, 100%) were compatible with the phylogeny, whereas this was the case for only 53 of 66 (80.3%) of 7-gene STs: 12 STs were polyphyletic and one was paraphyletic.

The global distribution of *C. diphtheriae* sublineages, apparent from the current sample of 602 genomes, revealed a strong phylogeographic pattern with most sublineages being typically restricted to one or a few adjacent countries (**Figure 3**). For example, SL8 was restricted to Europe, SL405 to India and SL40 to Indonesia. A notable exception was SL32, which was observed in Australia, Italy and the UK. SL5 was isolated in Europe except one isolate from Australia.

**Figure 3.**
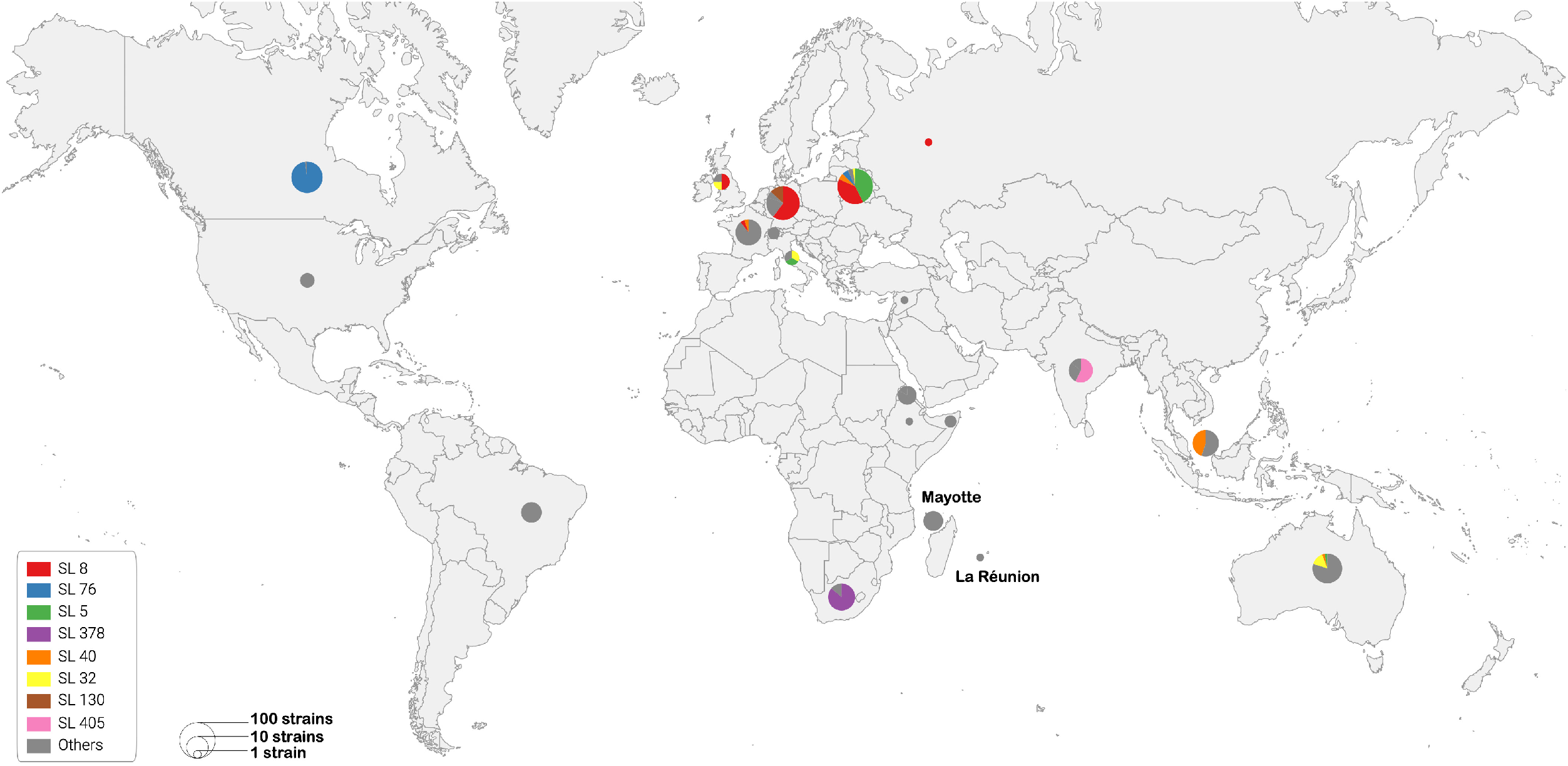
Geographic distribution of the most frequent genome-sequenced *C. diphtheriae* sublineages. The geographic origins of *C. diphtheriae* sublineages is represented by circles, the size of which represents the number of genomes per country (see key), considering the study dataset of 440 genomes as well as the 162 additional recently published genomes. The eight main sublineages are represented by colors of the pie charts sectors (see key); the other sublineages are grouped in the ‘others’ category and represented in grey.

### Frequently sampled sublineages of *C. diphtheriae*

The most frequent sublineages in our dataset included 7-gene MLST genotypes ST5, ST8, ST76 and ST378; they were labeled as SL5, SL8, SL76 and SL378, respectively, by our inheritance script. These sublineages were previously reported from diphtheria outbreaks. SL378 comprised 19 *tox*-positive isolates (except one tox-negative) recovered during a 20151016 outbreak in the KwaZulu province of South Africa [25]. Sublineage SL76 comprised isolates that were all *tox*-negative and belonged to ST76 or ST746 (a single-locus variant of ST76). Forty-five isolates were drawn from the Vancouver 2015-2018 outbreak of *tox*-negative *C. diphtheriae* [21], and five ST76 isolates were recovered from Belarus between 2004 and 2014 (**Table S1**).

Sublineage SL5 comprised 39 *tox*-negative genomes belonging to ST5 (all except 6), ST728, ST741 and 4 genomes for which the ST cannot be defined because they lack a *leuA* allele call (**Figure 4A**). ST5 contributed to infections in Belarus from the mid-1990s until at least 2013 [16] and was also recovered from Italy, Australia (**Table S1**) and Russia [55][56]. As noted above, whereas a majority of ST5 genomes were *spuA* negative, a distinct subbranch within SL5 possessed the biovar Gravis marker *spuA*. The phylogenetic analysis of SL5 genomes suggests initial acquisition of *spuA* in the ancestral branch of SL5 (after ~ year 1834), followed by its loss in a ST5 subbranch before 1955 (**Figure 1; Figure 4A**).

**Figure 4.**
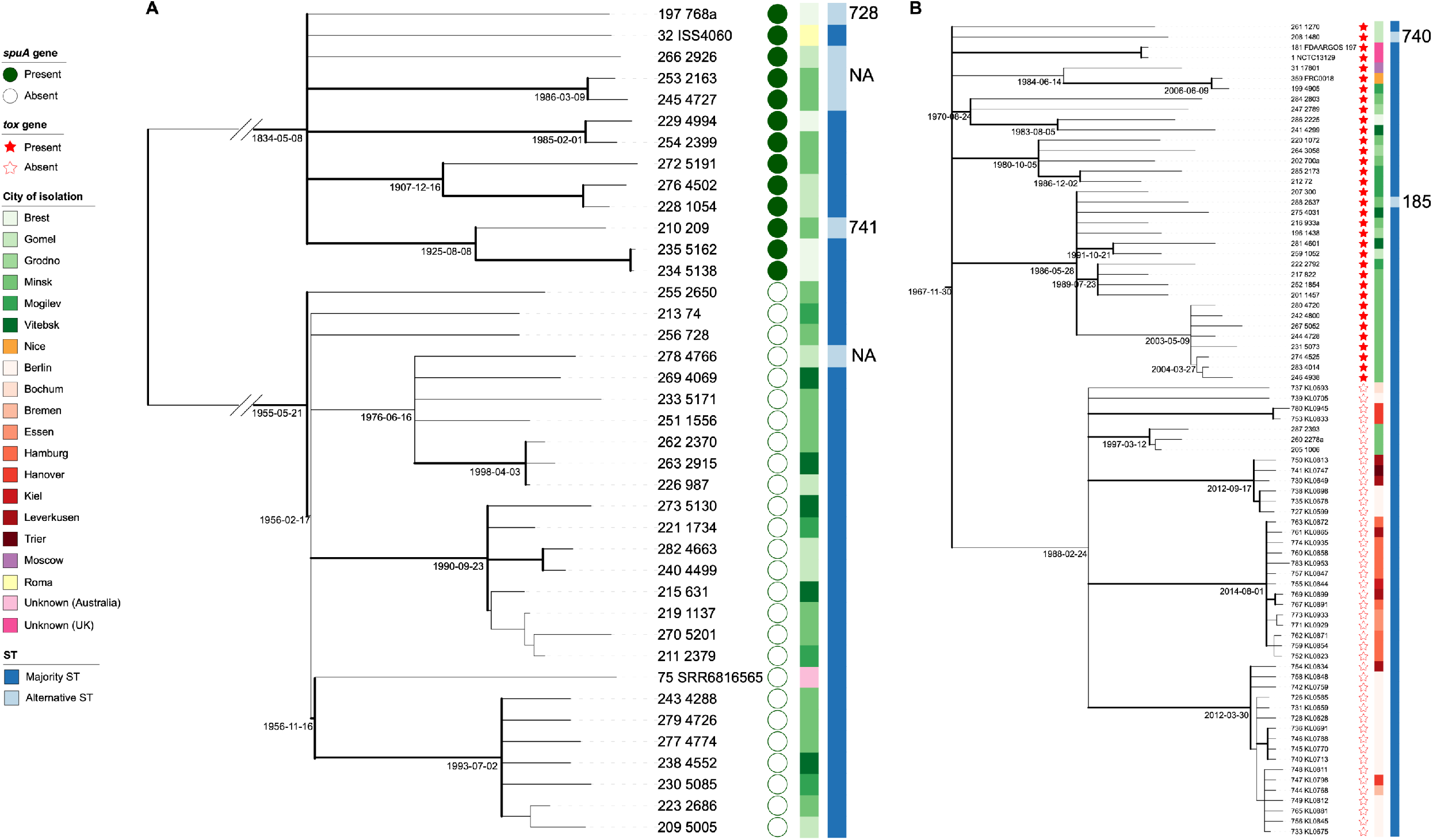
Time-scaled phylogenetic trees of SL5 and SL8 sublineages. Panel A: Sublineage 5; Panel B: sublineage 8. The phylogenies were obtained based on the LSD method using IQ-TREE v2.0.6 (see Methods); multifurcations were created when estimated node dates were conflicting with topology. The genomes are labeled with their BIGSdb identifier followed after a space by their original strain name. Estimated dates of major nodes are given below the left end of the branch above the node. Thick branches have a bootstrap support value greater than 70%. In panel A, circles represent the presence (green filled) or absence (empty) of the *spuA* gene. All strains were *tox*-negative (see main Figure 1). In panel B, the red stars the presence (red filled) or absence (empty) of the *tox* gene. All strains were *spuA* positive (see main Figure 1). In both panels, the central color strip indicates the isolates’ city of isolation, while the rightmost color strip shows the ST of isolates: majority STs in dark blue (ST5 for SL5 and ST8 for SL8) or alternative STs in light blue; the latter are labelled on the right; NA: not applicable (lack of *leuA* allele call in four SL5 genomes).

SL8 was isolated from cases of the large ex-Soviet Union outbreak [35] and the genomes of this sublineage were isolated over a period of 20 years from Belarus, Germany, Russia and the UK (**Table S1**) [35][29][16][20][57]. Strikingly, SL8 isolates were grouped into two subbranches, which were either *tox*-positive (subbranch A) or tox-negative (subbranch B; **Figure 1; Figure 4B**). Phylogenetic sister groups of the ST8 sublineage were ST123, ST451 and ST441, which comprised only *tox*-negative isolates. These data suggest a scenario of acquisition of the *tox* gene in the branch leading to the SL8 ancestor (before ~1967), and its subsequent loss in subbranch B (before ~1988).

### cgMLST variation within clusters or outbreak sets *versus* sporadic isolates and comparison with whole-genome single nucleotide polymorphisms

We aimed to assess the cgMLST variation within sets of isolates defined as epidemiologically related, *i.e*., described as case clusters or outbreaks (hereafter, clusters) based on their provenance information. Fifteen such sets were collated, including four clusters identified by the French national surveillance system and sequenced for the purposes of this study (**Table 1**). The 15 clusters of infections were caused by diverse strains, which were largely scattered across the phylogeny of *C. diphtheriae* (**Figure 1**). The distribution of allelic mismatches was clearly distinct for clusters *versus* non-related sets of isolates (**Figure 5**), and whereas the number of allelic mismatches observed within clusters ranged from none to 24, allelic variation among non-related isolates typically had more than 1000 mismatches.

**Figure 5.**
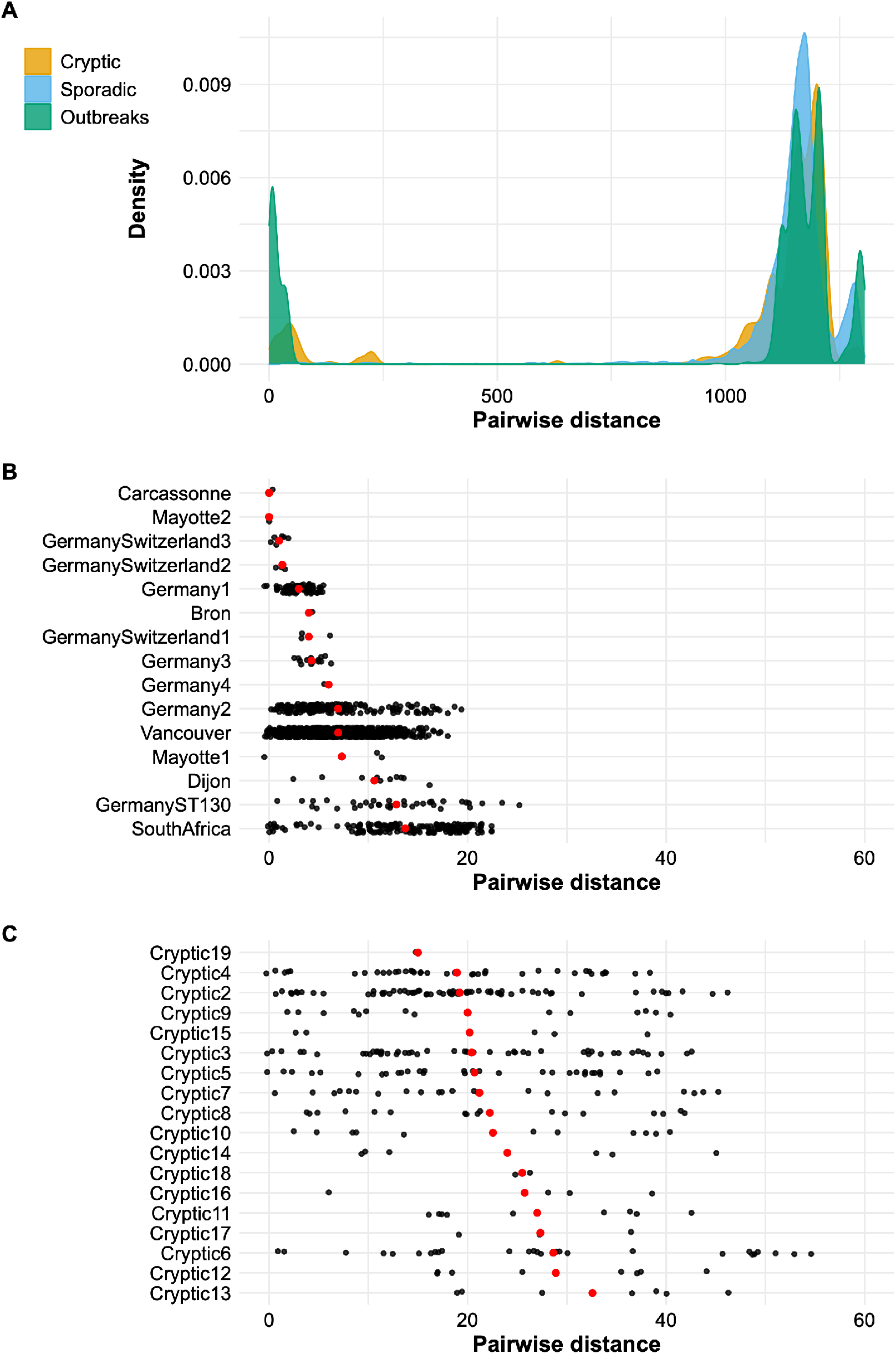
Density distribution of the pairwise cgMLST allelic distances among sporadic, cryptic clusters or outbreak isolates. **A**: Distribution density of the pairwise distances among all pairs of isolates included in outbreak clusters (green), cryptic clusters (orange) and sporadic cases (blue). **B**: Distribution of the pairwise distances among isolates within each outbreak cluster. **C**: distribution of the pairwise distances among isolates within cryptic clusters; the strains belonging to each outbreak or cryptic cluster are given in **Table S1**. In B and C, each black dot represents a pairwise distance. Red dots represent the mean of each cluster.

To compare the discriminatory power of cgMLST with the expectedly higher resolution provided by whole genome SNPs, two outbreaks for which read sets were available were investigated. The pairwise number of SNPs versus cgMLST allelic mismatches showed a strong correlation and indicated that 30 SNPs are observed between pairs of genomes that differ by ~15 to 17 cgMLST mismatches (**Figure 6**). Minimum spanning tree analysis of the genomes showed a high concordance between both methods in inferring genetic relationships of closely-related genomes (**Figures S10 and S11**).

**Figure 6.**
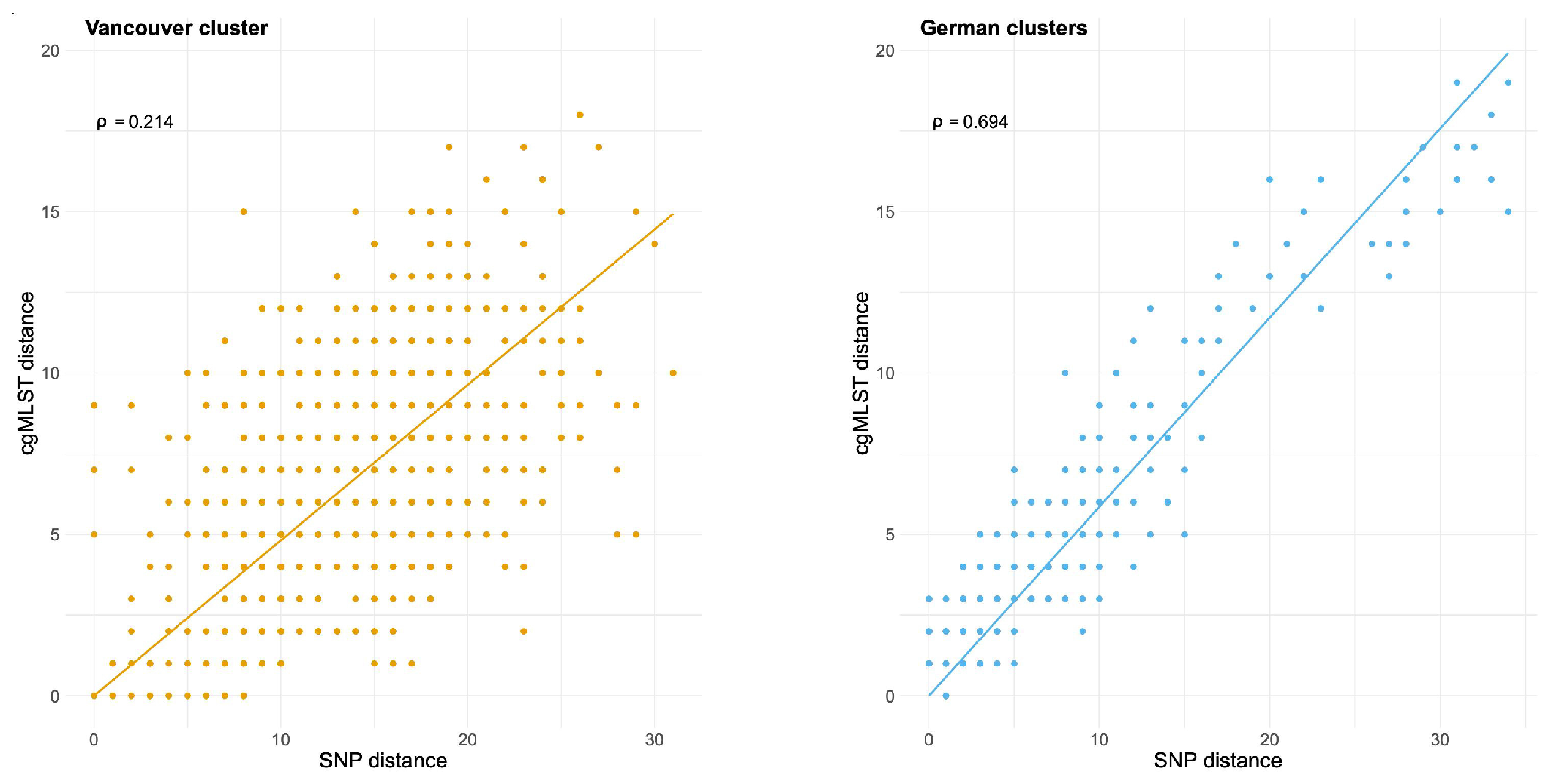
Relationships between the pairwise distances derived from whole-genome SNPs and cgMLST profiles, for two outbreak clusters. Each dot represents one pair of genomes. The line corresponds to the linear regression between the cgMLST **(Y-axis)** and SNP **(X-axis)** distances. The p value is the concordance correlation coefficient (ranging from −1 to 1). Left: Vancouver outbreak; Right: Germany clusters.

### cgMLST cluster classification uncovers cryptic transmission and cryptic duplicated cultures

We classified the 440 *C. diphtheriae* genomes into single linkage groups using a threshold of 25 cgMLST mismatches, corresponding to the maximum value observed so far for well described clusters. We define these groups as genetic clusters. This analysis enabled to detect 43 cryptic clusters, *i.e*., isolates belonging to genetic clusters but for which we have no evidence of epidemiological links. The cryptic clusters were genetically more heterogeneous than the epidemiological clusters (**Figure 5**). One cryptic cluster corresponded in fact to three distinct genome assemblies of the reference strain Park-Williams 8 (generally named PW8), which were deposited separately in public sequence databases (**Table S1**). These genomes differed by up to 20 loci among them, suggesting that multiple genetic differences exist among subcultures of this historical strain, isolated in 1896 and used for vaccine production. Likewise, there were four separately submitted genome sequences of strain C7beta (**Table S1**), but these showed only <3 cgMLST distinct alleles. Cryptic cluster 8 included NCTC3529 and NCTC5011 (which are synonyms) and strain c21 from India, which differed from the two others by 22 loci.

Of the remaining 40 cryptic clusters, 33 were single-country clusters and may correspond to local chains of transmission that were previously undetected, or just unreported to our knowledge. Only five cryptic clusters included isolates from at least two countries. Cluster 1 (as numbered in **Table S1**) corresponded to isolates from Belarus and the UK isolated between 1996 and 2004, and belong to the SL8 sublineage involved in the large ex-Soviet Union countries outbreak. Cryptic cluster 26 included two isolates from year 2008, one from Belarus and one from France, and the French patient had a history of travel to Russia; this cluster also belonged to SL8. Cryptic cluster 5 involved two isolates, one from Berlin in 2017 and one from Rio de Janeiro in 1999. Cryptic cluster 15 involved isolates from 2015 and 2016 in Australia and Germany. Cryptic 38 included isolates from France, Germany and Australia, 2013-2015. Cryptic cluster 3 involved two strains from Brazil, which were isolated in 1981 and 1993. Finally, we also revisited the clades defined in a recent study of the isolates from the large diphtheria outbreak in Yemen and found 13 groups, one of which (genetic cluster 217) corresponded to the predominant Yemen sublineage A.1.1, which was estimated to have originated in March 2015 [9].

## Discussion

*C. diphtheriae* was previously described as a genetically heterogeneous species [58][11, 18]. Here we compared all available genomic sequences and defined *C. diphtheriae* sublineages, showing that they are numerous and still largely under-sampled. Sublineages are separated by large genetic distances (2.4% on average), which indicates that *C. diphtheriae* is an old species that may have existed since several million years [59]. The relatively rapid evolutionary rate, which was previously estimated at ~1.10^-6^ substitutions per site year^-1^ within several *C. diphtheriae* sublineages [16, 60], and the frequent homologous recombination, which drives evolutionary divergence approximately 5 times faster than mutation [11], both contribute to the diversification of extant *C. diphtheriae* strains. This high level of genetic variation and fast diversification underline the need for a standard definition of *C. diphtheriae* strains and sublineages based on a precise genome-based genotyping approach. So far, this lack has limited our understanding of the patterns of geographical spread and of the phenotypic and clinical differences among *C. diphtheriae* sublineages.

Here, we developed a cgMLST genotyping approach, designed to be widely applicable across *C. diphtheriae* and its closely related species *C. rouxii* and *C. belfantii*. cgMLST is recognized as a highly reproducible and discriminatory approach applicable to molecular epidemiology and population biology questions [24, 61, 62]. We proposed a definition of sublineages that considers the population structure, and ensured backward compatibility with MLST identifiers through their inheritance onto cgMLST-based sublineages. In the future, novel sublineage identifiers will not be matched to MLST for practical reasons, and we propose that the cgMLST nomenclature should be prioritized over MLST. We integrated into the nomenclature, all publicly available genomes as of April 2021. The resulting genomic library is open to future community submissions and curator contributions.

Although most sublineages appear geographically restricted, sampling is currently very limited and the current picture is strongly biased by a focus on local transmission and outbreaks. It is striking that, excluding outbreaks, no *C. diphtheriae* sublineage is numerically predominant, suggesting a lack of globally successful clonal expansions. This contrasts with the situation observed in multiple bacterial pathogens where specific sublineages have come to predominate [61, 62]. The population dynamics of *C. diphtheriae* and its ecological drivers remains largely to be understood, but the evidence so far points to a large reservoir of diversity, strong phylogeographic structure and a lack of rapid emergence and spread of predominant sublineages. Asymptomatic carriage of *C. diphtheriae* in human populations may play an important role in maintaining the large genetic pool of *C. diphtheriae*.

We used a threshold of 25 allelic mismatches to define genetic clusters. This threshold captured all documented single-strain outbreak or case clusters. However, looser transmission chains of epidemiological relevance may exist and the threshold should be considered as indicative. Cryptic cluster analysis illustrates the potential value of cgMLST for uncovering previously hidden links between *C. diphtheriae* infection cases. Using our definition, we noted that high diversity of genetic clusters exists within single outbreaks of classical diphtheria, such as in the ex-Soviet Union or in Yemen. These observations underline the diversity of transmission chains that can occur simultaneously during such events, consistent with local reservoirs of diversity and silent circulation of *C. diphtheriae* until disease protection vanishes following a decrease in vaccination coverage. Strain heterogeneity during outbreaks may imply variations in toxin production or antimicrobial sensitivity, which could be highly relevant in terms of patient care and infection control priorities. The variation observed among subcultures of strain PW8 interrogates as to the possible impact on vaccine effectiveness, of the use of distinct variants of this strain in vaccine formulations [63].

The 1,305 cgMLST loci cover only approximately half of the entire genome length of a typical *C. diphtheriae*. Hence, the cgMLST has an inherent limitation in capturing genetic variation. Because our main goal was classification and nomenclature, we filtered out multiple genes to prioritize genotyping robustness and call rate over strain typing resolution power. Still, our comparison of cgMLST and whole genome SNP variation within described outbreaks shows high concordance between these methods, while illustrating that SNP calling will provide some additional resolution when needed. However, this more resolutive approach does not allow straightforward comparison of the genotypes sampled across different studies, and the two approaches should therefore be viewed as complementary.

Our phylogenetic analyses show that *C. diphtheriae* genomes can acquire and lose, at small evolutionary timescales, gene regions that underlie toxigenicity and biovar. The phyletic pattern of the *spuA* gene presence suggests that both losses and gains may be selectively advantageous given the circumstances. Likewise, the *tox* gene has been acquired or lost a large number of times in the population history of *C. diphtheriae*, and some of these events were traced here to a few years or decades back. Together with antimicrobial resistance and other genes acquisition and loss [58][11, 64], these evolutionary patterns paint a picture of *C. diphtheriae* as a highly dynamic species in which sublineages may evolve at epidemiological timescales with respect to their medically important characteristics. This underlines the need to better understand the drivers of these changes and to monitor the emergence and spread of genotypes of concern, *e.g*. which would combine multidrug resistant and toxigenic properties. Here, we did not look in details at the accessory genome, and future genotype-phenotype relationships studies should be carried out to decipher the impact of *C. diphtheriae* diversity on the heterogeneity of clinical phenotypes.

Important evolutionary and epidemiological questions about *C. diphtheriae* remain unanswered, including the likelihood for *tox*-negative strains to turn positive during infection or short-term transmission. Asymptomatic carriage is also poorly understood, and whether *C. diphtheriae* has evolved to partially escape vaccine-induced immunity in the context of long-standing vaccination [63] is currently unknown. This work provides a framework to harmonize knowledge on the infra-specific diversity of *C. diphtheriae*. The use of a unified approach to define *C. diphtheriae* strains should facilitate advances of our knowledge of the evolutionary and transmission dynamics of *C. diphtheriae* and the identification of strains with particular clinical properties or emerging behavior.

## Nucleotide sequence accession numbers

Newly sequenced isolates have been deposited to the European Nucleotide Archive (from ERS6637895 to ERS6637897).

## Content category

## Abbreviations

cgMLST: core genome Multilocus Sequence Typing

## Acknowledgements

We thank Melody Dazas for assistance in the early steps of the project and Annick Carmi-Leroy, Annie Landier, Nathalie Armatys and Virginie Passet for technical assistance with the microbiological characterization and sequencing of the *Corynebacterium diphtheriae* strains from the National Reference Center. We thank Vincent Enouf and the P2M core facility of Institut Pasteur for genomic sequencing. This work used the computational and storage services (TARS cluster) provided by the IT department at the Institut Pasteur, Paris.

## Ethical approval statement

To conduct the research, we used bacterial strains, which are not considered human samples. Accordingly, this research was not considered human research and is out of the scope of the decree n° 2016-1537 of November 16, 2016 implementing law n° 2012-300 of March 5, 2012 on research involving human subjects. Therefore, no ethics approval was needed and no informed consent was required.

All French bacteriological samples were collected in the frame of the French national diphtheria surveillance and are collected, coded, shipped, managed and analyzed according to the French National Reference Center protocols that received approval by French competent body CNIL (Commission Nationale Informatique et Libertés; approval number: 1474671). Other strains were obtained from culture collections.

The research conformed to the principles of the Helsinki Declaration.

## Authors license statement

This research was funded, in whole or in part, by Institut Pasteur and Santé publique France. For the purpose of open access, the authors have applied a CC-BY public copyright license to any Author Manuscript version arising from this submission.

## Author contributions

S.B. conceived, designed and coordinated the study. E.B. and S.B. coordinated the microbiological cultures of the isolates and their biochemical and molecular characterizations. J.G., M.H., A.C. and S.B. analyzed the genomic data. J.T. reviewed the clinical source data of the isolates collection. S.B. and J.G. wrote the initial version of the manuscript. All authors provided input to the manuscript and reviewed the final version.

## References

1. Burkowski A. Diphtheria and its Etiological Agents. In: Corynebacterium diphtheriae and Related Toxigenic Species. Andreas Burkowski. pp. 1–10.

2. Zasada AA, Zaleska M, Podlasin RB, Seferyńska I. The first case of septicemia due to nontoxigenic Corynebacterium diphtheriae in Poland: case report. Ann Clin Microbiol Antimicrob 2005;4:8.

3. Sharma NC, Efstratiou A, Mokrousov I, Mutreja A, Das B, et al. Diphtheria. Nat Rev Dis Primer 2019;5:81.

4. ECDC. European Centre for Disease Prevention and Control. Diphtheria. Annual epidemiological report for 2018. https://www.ecdc.europa.eu/sites/default/files/documents/diphtheria-annual-epidemiological-report-2018.pdf (2021).

5. Meinel DM, Kuehl R, Zbinden R, Boskova V, Garzoni C, et al. Outbreak investigation for toxigenic Corynebacterium diphtheriae wound infections in refugees from Northeast Africa and Syria in Switzerland and Germany by whole genome sequencing. Clin Microbiol Infect Off Publ Eur Soc Clin Microbiol Infect Dis 2016;22:1003.e1–1003.e8.

6. Scheifer C, Rolland-Debord C, Badell E, Reibel F, Aubry A, et al. Re-emergence of Corynebacterium diphtheriae. Med Mal Infect 2019;49:463–466.

7. Dittmann S, Wharton M, Vitek C, Ciotti M, Galazka A, et al. Successful control of epidemic diphtheria in the states of the Former Union of Soviet Socialist Republics: lessons learned. J Infect Dis 2000;181 Suppl 1:S10–22.

8. Polonsky JA, Ivey M, Mazhar MKA, Rahman Z, le Polain de Waroux O, et al. Epidemiological, clinical, and public health response characteristics of a large outbreak of diphtheria among the Rohingya population in Cox’s Bazar, Bangladesh, 2017 to 2019: A retrospective study. PLoS Med 2021;18:e1003587.

9. Badell E, Alharazi A, Criscuolo A, Almoayed KAA, Lefrancq N, et al. Ongoing diphtheria outbreak in Yemen: a cross-sectional and genomic epidemiology study. Lancet Microbe;0. Epub ahead of print 26 May 2021. DOI: 10.1016/S2666-5247(21)00094-X.

10. Sangal V, Burkovski A, Hunt AC, Edwards B, Blom J, et al. A lack of genetic basis for biovar differentiation in clinically important Corynebacterium diphtheriae from whole genome sequencing. Infect Genet Evol J Mol Epidemiol Evol Genet Infect Dis 2014;21:54–57.

11. Hennart M, Panunzi LG, Rodrigues C, Gaday Q, Baines SL, et al. Population genomics and antimicrobial resistance in Corynebacterium diphtheriae. Genome Med 2020;12:107.

12. Dazas M, Badell E, Carmi-Leroy A, Criscuolo A, Brisse S. Taxonomic status of Corynebacterium diphtheriae biovar Belfanti and proposal of Corynebacterium belfantii sp. nov. Int J Syst Evol Microbiol 2018;68:3826–3831.

13. Badell E, Hennart M, Rodrigues C, Passet V, Dazas M, et al. Corynebacterium rouxii sp. nov., a novel member of the diphtheriae species complex. Res Microbiol. Epub ahead of print 28 February 2020. DOI: 10.1016/j.resmic.2020.02.003.

14. Bolt F, Cassiday P, Tondella ML, Dezoysa A, Efstratiou A, et al. Multilocus sequence typing identifies evidence for recombination and two distinct lineages of Corynebacterium diphtheriae. J Clin Microbiol 2010;48:4177–4185.

15. Grimont PAD, Grimont F, Efstratiou A, De Zoysa A, Mazurova I, et al. International nomenclature for Corynebacterium diphtheriae ribotypes. Res Microbiol 2004;155:162–166.

16. Grosse-Kock S, Kolodkina V, Schwalbe EC, Blom J, Burkovski A, et al. Genomic analysis of endemic clones of toxigenic and non-toxigenic Corynebacterium diphtheriae in Belarus during and after the major epidemic in 1990s. BMC Genomics 2017;18:873.

17. Timms VJ, Nguyen T, Crighton T, Yuen M, Sintchenko V. Genome-wide comparison of Corynebacterium diphtheriae isolates from Australia identifies differences in the Pan-genomes between respiratory and cutaneous strains. BMC Genomics 2018;19:869.

18. Seth-Smith HMB, Egli A. Whole Genome Sequencing for Surveillance of Diphtheria in Low Incidence Settings. Front Public Health 2019;7:235.

19. Berger A, Dangel A, Schober T, Schmidbauer B, Konrad R, et al. Whole genome sequencing suggests transmission of Corynebacterium diphtheriae-caused cutaneous diphtheria in two siblings, Germany, 2018. Euro Surveill Bull Eur Sur Mal Transm Eur Commun Dis Bull;24. Epub ahead of print January 2019. DOI: 10.2807/1560-7917.ES.2019.24.2.1800683.

20. Dangel A, Berger A, Konrad R, Bischoff H, Sing A. Geographically Diverse Clusters of Nontoxigenic Corynebacterium diphtheriae Infection, Germany, 2016-2017. Emerg Infect Dis 2018;24:1239–1245.

21. Chorlton SD, Ritchie G, Lawson T, Romney MG, Lowe CF. Whole-genome sequencing of Corynebacterium diphtheriae isolates recovered from an inner-city population demonstrates the predominance of a single molecular strain. J Clin Microbiol. Epub ahead of print 20 November 2019. DOI: 10.1128/JCM.01651-19.

22. Pivot D, Fanton A, Badell-Ocando E, Benouachkou M, Astruc K, et al. Carriage of a single strain of non-toxigenic Corynebacterium diphtheriae biovar Belfanti (Corynebacterium belfantii) in four patients with cystic fibrosis. J Clin Microbiol. Epub ahead of print 27 February 2019. DOI: 10.1128/JCM.00042-19.

23. Xiaoli L, Benoliel E, Peng Y, Aneke J, Cassiday PK, et al. Genomic epidemiology of nontoxigenic Corynebacterium diphtheriae from King County, Washington State, USA between July 2018 and May 2019. Microb Genomics;6. Epub ahead of print December 2020. DOI: 10.1099/mgen.0.000467.

24. Maiden MC, van Rensburg MJ, Bray JE, Earle SG, Ford SA, et al. MLST revisited: the gene-by-gene approach to bacterial genomics. Nat Rev Microbiol 2013;11:728–36.

25. du Plessis M, Wolter N, Allam M, de Gouveia L, Moosa F, et al. Molecular Characterization of Corynebacterium diphtheriae Outbreak Isolates, South Africa, March-June 2015. Emerg Infect Dis 2017;23:1308–1315.

26. Pivot D, Fanton A, Badell-Ocando E, Benouachkou M, Astruc K, et al. Carriage of a Single Strain of Nontoxigenic Corynebacterium diphtheriae bv. Belfanti (Corynebacterium belfantii) in Four Patients with Cystic Fibrosis. J Clin Microbiol;57. Epub ahead of print May 2019. DOI: 10.1128/JCM.00042-19.

27. Schaeffer J, Huhulescu S, Stoeger A, Allerberger F, Ruppitsch W. Assessing the Genetic Diversity of Austrian Corynebacterium diphtheriae Clinical Isolates, 2011-2019. J Clin Microbiol. Epub ahead of print 2 December 2020. DOI: 10.1128/JCM.02529-20.

28. Hoefer A, Pampaka D, Herrera-León S, Peiró S, Varona S, et al. Molecular and epidemiological characterisation of toxigenic and non-toxigenic C. diphtheriae, C. belfantii and C. ulcerans isolates identified in Spain from 2014 to 2019. J Clin Microbiol. Epub ahead of print 9 December 2020. DOI: 10.1128/JCM.02410-20.

29. Will RC, Ramamurthy T, Sharma NC, Veeraraghavan B, Sangal L, et al. Spatiotemporal persistence of multiple, diverse clades and toxins of Corynebacterium diphtheriae. Nat Commun 2021;12:1500.

30. Criscuolo A, Brisse S. AlienTrimmer: A tool to quickly and accurately trim off multiple short contaminant sequences from high-throughput sequencing reads. Genomics 2013;10.1016/j.ygeno.2013.07.011.

31. Liu Y, Schröder J, Schmidt B. Musket: a multistage k-mer spectrum-based error corrector for Illumina sequence data. Bioinforma Oxf Engl 2013;29:308–315.

32. Magoč T, Salzberg SL. FLASH: fast length adjustment of short reads to improve genome assemblies. Bioinformatics 2011;27:2957–2963.

33. Bankevich A, Nurk S, Antipov D, Gurevich AA, Dvorkin M, et al. SPAdes: a new genome assembly algorithm and its applications to single-cell sequencing. J Comput Biol 2012;19:455–77.

34. Guglielmini J, Bourhy P, Schiettekatte O, Zinini F, Brisse S, et al. Genus-wide Leptospira core genome multilocus sequence typing for strain taxonomy and global surveillance. PLoS Negl Trop Dis 2019;13:e0007374.

35. Cerdeño-Tárraga AM, Efstratiou A, Dover LG, Holden MTG, Pallen M, et al. The complete genome sequence and analysis of Corynebacterium diphtheriae NCTC13129. Nucleic Acids Res 2003;31:6516–6523.

36. Wozniak M, Wong L, Tiuryn J. eCAMBer: efficient support for large-scale comparative analysis of multiple bacterial strains. BMCBioinformatics 2014;15:65.

37. Hyatt D, Chen G-L, LoCascio PF, Land ML, Larimer FW, et al. Prodigal: prokaryotic gene recognition and translation initiation site identification. BMC Bioinformatics 2010;11:119.

38. Touchon M, Hoede C, Tenaillon O, Barbe V, Baeriswyl S, et al. Organised genome dynamics in the Escherichia coli species results in highly diverse adaptive paths. PLoS Genet 2009;5:e1000344.

39. Kent WJ. BLAT—The BLAST-Like Alignment Tool. Genome Res 2002;12:656–664.

40. Jolley KA, Bliss CM, Bennett JS, Bratcher HB, Brehony C, et al. Ribosomal multilocus sequence typing: universal characterization of bacteria from domain to strain. Microbiology 2012;158:1005–15.

41. Tagini F, Pillonel T, Croxatto A, Bertelli C, Koutsokera A, et al. Distinct Genomic Features Characterize Two Clades of Corynebacterium diphtheriae: Proposal of Corynebacterium diphtheriae Subsp. diphtheriae Subsp. nov. and Corynebacterium diphtheriae Subsp. lausannense Subsp. nov. Front Microbiol 2018;9:1743.

42. Jolley KA, Maiden MCJ. BIGSdb: Scalable analysis of bacterial genome variation at the population level. BMC Bioinformatics 2010;11:595.

43. Dangel A, Berger A, Konrad R, Sing A. NGS-based phylogeny of diphtheria-related pathogenicity factors in different Corynebacterium spp. implies species-specific virulence transmission. BMC Microbiol 2019;19:28.

44. Criscuolo A. A fast alignment-free bioinformatics procedure to infer accurate distance-based phylogenetic trees from genome assemblies. Res Ideas Outcomes 2019;5:e36178.

45. Didelot X, Wilson DJ. ClonalFrameML: efficient inference of recombination in whole bacterial genomes. PLoS Comput Biol 2015;11:e1004041.

46. Katoh K, Standley DM. MAFFT multiple sequence alignment software version 7: improvements in performance and usability. Mol Biol Evol 2013;30:772–780.

47. Bruen TC, Philippe H, Bryant D. A simple and robust statistical test for detecting the presence of recombination. Genetics 2006;172:2665–2681.

48. Rousseeuw PJ. Silhouettes: A graphical aid to the interpretation and validation of cluster analysis. J Comput Appl Math 1987;20:53–65.

49. To T-H, Jung M, Lycett S, Gascuel O. Fast Dating Using Least-Squares Criteria and Algorithms. Syst Biol 2016;65:82–97.

50. Minh BQ, Schmidt HA, Chernomor O, Schrempf D, Woodhams MD, et al. IQ-TREE 2: New Models and Efficient Methods for Phylogenetic Inference in the Genomic Era. Mol Biol Evol 2020;37:1530–1534.

51. Seemann T. snippy: fast bacterial variant calling from NGS reads. https://github.com/tseemann/snippy (2015).

52. Lomolino MV. Ecology’s most general, yet protean 1 pattern: the species-area relationship. J Biogeogr 2000;27:17–26.

53. Badell E, Hennart M, Rodrigues C, Passet V, Dazas M, et al. Description of Corynebacterium rouxii sp. nov., a novel member of the diphtheriae species complex. bioRxiv 2019;855692.

54. Santos AS, Ramos RT, Silva A, Hirata R, Mattos-Guaraldi AL, et al. Searching whole genome sequences for biochemical identification features of emerging and reemerging pathogenic Corynebacterium species. Funct Integr Genomics 2018;18:593–610.

55. Bolt F, Cassiday P, Tondella ML, Dezoysa A, Efstratiou A, et al. Multilocus sequence typing identifies evidence for recombination and two distinct lineages of Corynebacterium diphtheriae. J Clin Microbiol 2010;48:4177–4185.

56. Borisova OI, Mazurova IK, Chagina IA, Pimenova AS, Donskikh EE, et al. [Multilocus sequencing of Corynebacterium diphtheriae strains isolated in Russia in 2002 - 2012]. Zh Mikrobiol Epidemiol Immunobiol 2013;17–23.

57. Farfour E, Badell E, Zasada A, Hotzel H, Tomaso H, et al. Characterization and comparison of invasive Corynebacterium diphtheriae isolates from France and Poland. J Clin Microbiol 2012;50:173–175.

58. Sangal V, Hoskisson PA. Evolution, epidemiology and diversity of Corynebacterium diphtheriae: New perspectives on an old foe. Infect Genet Evol J Mol Epidemiol Evol Genet Infect Dis 2016;43:364–370.

59. Achtman M, Wagner M. Microbial diversity and the genetic nature of microbial species. Nat Rev Microbiol 2008;6:431–40.

60. Badell E, Alharazi A, Criscuolo A, Group TN diphtheria outbreak working, Lefrancq N, et al. Epidemiological, clinical and genomic insights into the ongoing diphtheria outbreak in Yemen. medRxiv 2020;2020.07.21.20159186.

61. Alikhan N-F, Zhou Z, Sergeant MJ, Achtman M. A genomic overview of the population structure of Salmonella. PLoS Genet 2018;14:e1007261.

62. Moura A, Criscuolo A, Pouseele H, Maury MM, Leclercq A, et al. Whole genome-based population biology and epidemiological surveillance of Listeria monocytogenes. Nat Microbiol 2016;2:16185.

63. Möller J, Kraner M, Sonnewald U, Sangal V, Tittlbach H, et al. Proteomics of diphtheria toxoid vaccines reveals multiple proteins that are immunogenic and may contribute to protection of humans against Corynebacterium diphtheriae. Vaccine 2019;37:3061–3070.

64. Trost E, Blom J, Soares S de C, Huang I-H, Al-Dilaimi A, et al. Pangenomic study of Corynebacterium diphtheriae that provides insights into the genomic diversity of pathogenic isolates from cases of classical diphtheria, endocarditis, and pneumonia. J Bacteriol 2012;194:3199–3215.

